# LipidFinder 2.0: advanced informatics pipeline for lipidomics discovery applications

**DOI:** 10.1101/2020.08.16.250878

**Authors:** Jorge Alvarez-Jarreta, Patricia R.S. Rodrigues, Eoin Fahy, Anne O’Connor, Anna Price, Caroline Gaud, Simon Andrews, Paul Benton, Gary Siuzdak, Jade I. Hawksworth, Maria Valdivia-Garcia, Stuart M. Allen, Valerie B. O’Donnell

**Affiliations:** Systems Immunity Research Institute, School of Medicine, Cardiff University, CF14 4XN, UK; European Molecular Biology Laboratory, European Bioinformatics Institute (EMBL-EBI), Wellcome Genome Campus, Hinxton, CB10 1SD, UK; Department of Bioengineering, University of California, San Diego, California, USA; School of Biosciences, Cardiff University, CF10 3AX, UK; Bioinformatics, Babraham Institute, Cambridge, UK; The Scripps Research Institute, Center for Metabolomics, CA 92037; School of Computer Science and Informatics, Cardiff University, CF24 3AA, UK

## Abstract

We present LipidFinder 2.0, incorporating four new modules that apply artefact filters, remove lipid and contaminant stacks, in-source fragments and salt clusters, and a new isotope deletion method which is significantly more sensitive than available open-access alternatives. We also incorporate a novel false discovery rate (FDR) method, utilizing a target-decoy strategy, which allows users to assess data quality. A renewed lipid profiling method is introduced which searches three different databases from LIPID MAPS and returns bulk lipid structures only, and a lipid category scatter plot with color blind friendly pallet. An API interface with XCMS Online is made available on LipidFinder’s online version. We show using real data that LipidFinder 2.0 provides a significant improvement over non-lipid metabolite filtering and lipid profiling, compared to available tools.

**Availability:** LipidFinder 2.0 is freely available at https://github.com/ODonnell-Lipidomics/LipidFinder and http://lipidmaps.org/resources/tools/lipidfinder.

**Supplementary information:** Supplementary data are available at *Bioinformatics* online.

## 1 Introduction

Lipidomics describes the discovery and analysis of lipids (fats), which are essential molecules for life in all organisms (Wenk, 2005). However, it is hampered by the lack of specifically tailored informatics tools that effectively clean up raw datasets, which can contain huge numbers of artifacts (around 90-95% of initial signals) as described in detail in our Supplementary Information. Informatics tools used for lipidomics have been to a large extent designed for global metabolomics (and some are still focused primarily on this). The lack of specialized filters for lipidomics can have a negative effect on the output’s robustness since lipids have unique analytical challenges (Cappadona, et al., 2012; O'Donnell, et al., 2014). In 2017 we published a Python tool, LipidFinder, designed to be an additional stage of the lipidomics pipeline (O'Connor, et al., 2017). Specifically tailored for high-resolution LC/MS, LipidFinder processes the output of pre-processing tools, such as XCMS, to filter out common artefacts, including well known ESI contaminants and adducts, and remove background effects. LipidFinder was found to retain most reference lipids, improving the quality of lipidomic data. However, its output datasets still contained a significant level of artefacts, creating substantial problems for downstream statistical power. Thus, we developed new features and here we present LipidFinder 2.0, which integrates a new user-friendly interface to configure its parameters, additional filters and methods to improve the reliability and speed of the output.

A significant issue in identification is over-annotation where MS is used to assign a fully annotated structure. This has led to major mistakes in structural assignment (Bowden, et al., 2017; Liebisch, et al., 2015). This problem is unique to lipids due to the large numbers of isobaric ions. Currently open access lipidomics software don’t address this. Here we include an enhanced putative identification procedure that allows different degrees of profiling, returning only the bulk structure. We introduce a novel false discovery rate (FDR) method, using a target-decoy strategy. LipidFinder 2.0 supports as input datasets from popular pre-processing tools. We tested performance with real data, using XCMS-based lipidomics pipeline with/without LipidFinder and results are shown in Supplementary Results.

## 2 System and methods

The lipidomics analysis pipeline including LipidFinder 2.0 is outlined in Figure 1. Major changes have been introduced in its internal framework to improve usability, performance and reliability. These are explained in detail in the Supplementary Information and summarised here.

**Fig. 1.**
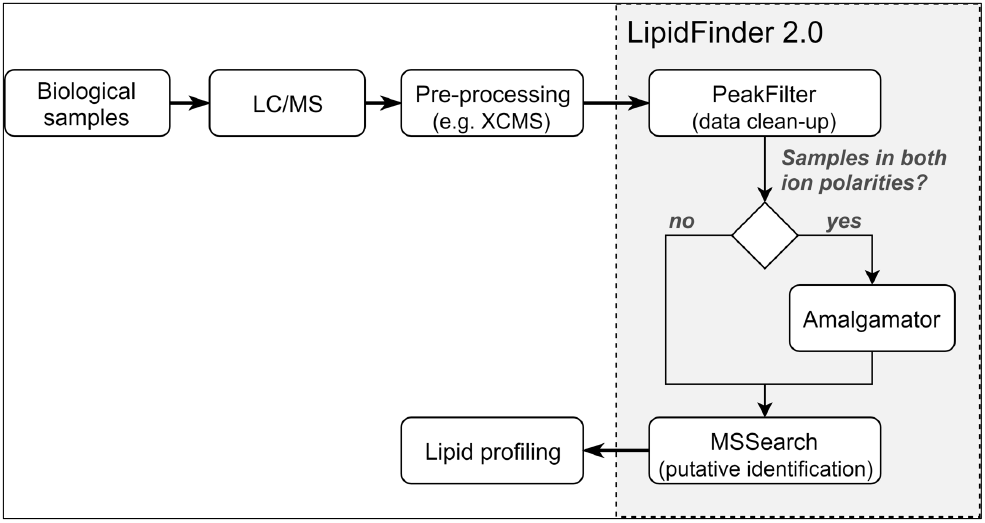
Common pipeline for untargeted lipidomics incorporating LipidFinder 2.0. It also shows LipidFinder’s new main workflow.

LipidFinder is designed to be integrated after the pre-processing stage in the popular XCMS-based data processing pipeline in lipidomics to improve the removal of artefacts, and to produce putative profiles enabling the statistical analysis of LC/MS datasets. New innovations are:

i. <u>A user-friendly configuration process for each module.</u> The configuration stage is designed for usability/adaptability for new or expert users, including a set of default settings customised for high-res LC/MS and a graphical user and command line interfaces. All widely used pre-processing tools can now be accommodated (Supplementary Methods).
ii. <u>Three new filters and a novel metric method added to the clean-up stage.</u> The new filters add in-source ion fragments, isotopes and salt clusters to the already broad list of common artefacts that LipidFinder targets and removes from MS datasets. Although isotope annotation can also be performed with CAMERA (included in XCMS), we found that its approach is not tailored optimally for lipidomics, thus we have implemented a new method with a more accurate intensity ratio check. Last, we have introduced a novel FDR method that serves as a metric to assess data quality. This is a new innovation not available in other lipidomics pipelines.
iii. <u>A comprehensive re-design of the profiling step to extend its applicability.</u> Here, the putative lipid profiling stage now returns lipid bulk structures rather than fully annotated, featuring three lipidomics databases from LIPID MAPS. It also returns lipid category scatter plots and fully annotated output files. All these are described in full (Supplementary Data).

## 3. Implementation

LipidFinder 2.0 is fully implemented in Python, supporting versions 2.7 and 3.3 or newer. The source code has been reorganized in a structure more similar to a Python library than a software tool, providing the scripts for the different stages shown in Section 2 and in Supplementary Results. The purpose is to encourage users with experience in programming to reuse LipidFinder’s filters or even entire stages to create their own tailored pipelines. Special attention was paid to *PeakFilter* to ensure it performs efficiently with large datasets. Also, as byproduct of our collaboration with LIPID MAPS (Fahy, et al., 2019), we developed an API that provides direct access to the databases, which has significantly reduced the time cost of *MS Search*. We have also linked LipidFinder on LIPID MAPS with XCMS Online to directly import pre-processed files. An analysis of the efficiency of the new implementation is shown in the Supplementary Results. We have produced the user manual in two formats: a PDF file and a Jupyter notebook. The latter converts it into an interactive cookbook where users can learn how to set up LipidFinder and how to use it.

The configuration files for each stage are now stored in JavaScript object notation (JSON) format, a readable text format that can be opened and modified by any text editor. Every data file involved in LipidFinder’s workflow, including those already provided with its source code, is saved either in comma-separated values (CSV) format (*PeakFilter*, *Amalgamator*) or Microsoft Excel spreadsheet (XLS and XLSX) format (*MS Search*), all of them widely supported by data handling applications.

Finally, we performed analysis with a biological dataset comprising lipids extracted from raw and peritoneal macrophages, in order to demonstrate the significant improvements in data quality and visualization achieved using LipidFinder 2.0, detailed in Supplementary Results.

## Funding

This project was supported by a Wellcome Trust grant for LIPID MAPS [203014/Z/16/Z]; and the European Research Council [LipidArrays]. VBO holds a Royal Society Wolfson Research Merit Award.

## Conflict of Interest

none declared.

## Supplementary Information

### Supplementary Introduction

Liquid chromatography-mass spectrometry (LC/MS) is an essential method for discovery and characterisation of lipids in biological samples. Pre-processing is a required step in the lipidomics pipeline, however most available informatics solutions are tailored for metabolomics and not specifically lipidomics and thus, they retain large numbers of artefactual ions in datasets. This leads to major challenges for statistical analysis and renders robust identification of novel lipids impossible.

Liquid chromatography coupled to mass spectrometry (LC/MS), is one of the most popular analysis methods used for profiling of lipids in biological samples. This method (untargeted lipidomics) is widely used due to its high coverage of molecules, aiming to support discovery of biomarkers related to development, disease and therapeutic response(Covey, 1986). However, depending on the experimental methodology and the tissue analysed, output datasets can contain up to tens of thousands of ions/peaks (features). For example, with human platelets, raw datasets can comprise close to 60 K features, of which it is estimated only around 4-5 K are real lipids(Slatter, et al., 2016). The rest are common artefacts, e.g. isotope peaks, electrospray ionisation (ESI) contaminant ions, salt clusters, in-source fragments and others. Also, a single lipid may be represented by many features. If these are not removed, two main problems result. First, novel lipids cannot be easily distinguished from non-lipids. Second, when comparing datasets for biomarker discovery, statistical power is reduced. Thus, new computational methods are urgently required to help researchers cope with this significant Big Data problem.

In lipidomics/metabolomics (Weckwerth, 2003), the common data processing pipeline starts with interrogation of biological samples using LC/MS. The output can be represented in a 3D chromatogram: mass-to-charge ratio (*m/z*) *vs* retention time *vs* signal intensity. It is essential to detect all “features” in the data, that is, every unique representation of *m/z* and retention time, and return these in a usable format for further analysis. Hence, the raw dataset needs to be “pre-processed”. For this, several tools have been implemented over the last 10-15 years: XCMS, MZmine and MetAlign are examples of open access options (Lommen and Kools, 2012; Pluskal, et al., 2010; Smith, et al., 2006). The last step of the pipeline is to match each feature to a known analyte, while also retaining unidentified ions for further analysis as potential new lipids. LC/MS is extremely useful for lipid discovery and comparative profiling. However, it is always recommended to follow LC/MS up with targeted methods such as tandem mass spectrometry (MS/MS), to validate the putative profiling returned by this “untargeted” lipidomics.

Pre-processing software tools for MS include two main methods devised to facilitate determining true lipid or metabolite ions: peak identification and peak alignment. The former aims to detect and extract every peak’s key information from the raw LC/MS data (e.g. mass centroid position and area under the curve). Peak alignment corrects for minor changes in retention time arising from fluctuations of environmental temperature and humidity, and health of chromatographic columns (Smith, et al., 2015; Zhang, et al., 2014). Insufficiently addressing alignment negatively impacts subsequent statistical analysis and identification of lipids/metabolites (Zhou, et al., 2012). Furthermore, some tools also include additional filters and methodologies to confirm that returned features correspond to known analytes. XCMS/CAMERA incorporates baseline and noise elimination, and an isotope and adduct detection method, as well as putative identification for features (Zhou, et al., 2012).

### Supplementary System and Methods

#### Input dataset and configuration process

In contrast to LipidFinder 1.0, which only worked with XCMS and SIEVE™ (ThermoFisher) datasets, LipidFinder 2.0 works with input files from widely used preprocessing tools. Here, the input file layout is far more flexible: a first column with a unique identifier per feature, *m/z* and retention time columns, and all sample measurement columns together are the only requirements. All columns except the identifier can be placed anywhere in the dataset. Also, the file can contain extra columns that can be retained in the output. The list of formats was extended so as well as comma-separated values (CSV), LipidFinder can read tab-separated values (TSV) and Microsoft Excel spreadsheet (XLS and XLSX) files.

Configuring LipidFinder 1.0 was considered challenging for inexperienced users, because it required editing a CSV file containing the parameter names, their description, restrictions and values. Thus, typos, incorrect data types and invalid values (due to threshold violations) were common mistakes during this step. To address this, a new configuration process with two interface options has been added to each stage. A graphical user interface (GUI) and a command-line interface (CLI) have been developed paying special attention to their usability to improve user interaction with the software. For example, we include an additional help text for parameters to provide support for new users and default values optimised for high-resolution LC/MS data. The resulting configuration file is still saved in a readable text format, so experienced users can edit it manually. We have also incorporated the functionality to transfer parameters between stages, that is, users can import the configuration file of a previous stage to copy the values of shared parameters, saving time and preventing errors during the configuration. Finally, LipidFinder 2.0 provides backward compatibility, so any configuration file from its previous version can be used as a foundation to generate the new ones.

#### PeakFilter

The first module in LipidFinder’s workflow is *PeakFilter*, designed to improve the quality of datasets through applying a series of corrective steps, outlined in Supplementary Figure 1. However, LipidFinder 1.0 still retained artefactual ions that need to be removed in order to ensure data quality. To this end, we include 4 new modules into *PeakFilter*, depicted in orange in Supplementary Figure 1, along with significantly improving the functionality of our Stack Removal module. These are described below.

**Supplementary Figure 1.**
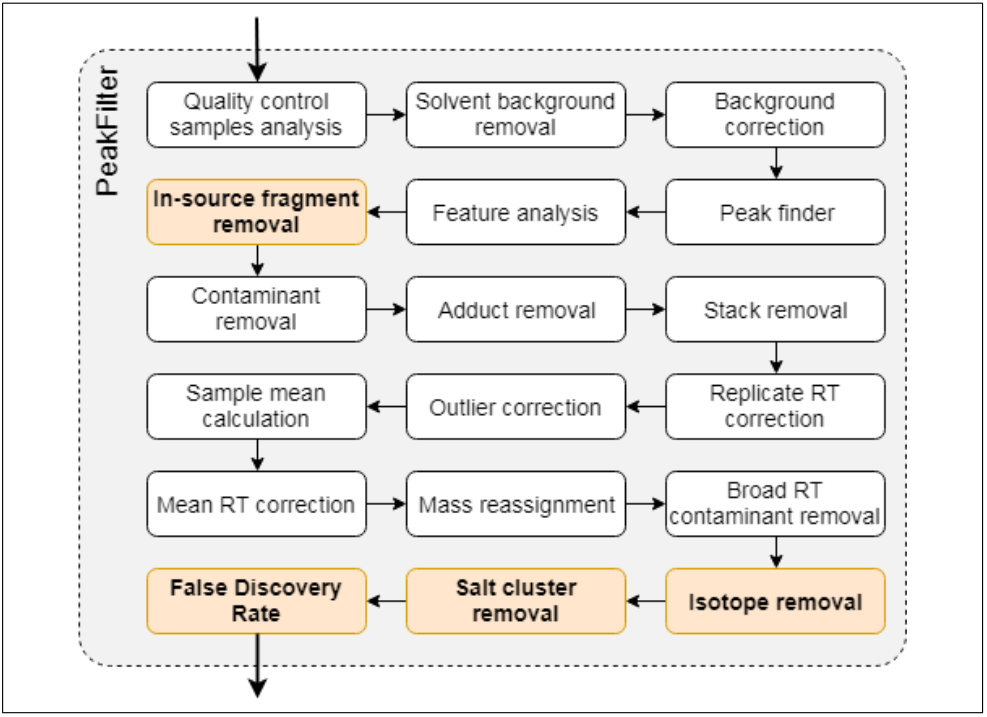
PeakFilter’s updated workflow. The 4 new modules are highlighted in orange.

##### 2.2.1 In-source fragmentation removal

In-source fragmentation is a common problem that arises in LC/MS where fragmentation of ions, to generate product ions, occurs in the source despite ESI being considered a soft ionization method(Gabelica and De Pauw, 2005). These product ions can be impossible to tell from real molecular ions and are misidentified using database searches, as they are isobaric with the precursor ions of real lipids (Xu, et al., 2015). To solve this issue, we developed a method that searches for common well-known lipid-related in-source ion fragments and removes them from the dataset. The most common forms of these are either **i)** fatty acyl ions, and related ketenes generated from phospholipids and detected in negative mode, and **ii)** neutral loss ions from the loss of common fatty acyls, headgroups, CO_2_ or H_2_O which can appear in either positive of negative mode. The default in-source fragments and common neutral losses targeted as well as the algorithm are provided in Supplementary Information (Appendix 1). The use of this module requires LC separation since default targets will induce the filter to remove free fatty acids unless clearly separated from larger lipids.

##### 2.2.2 Stack removal

LipidFinder 1.0 included a module to remove lipid and contaminant stacks. However, the implementation of this filter was too strict and some stacks remained in the resulting datasets. In this update we revisited the definition of both types of stacks to improve the accuracy of the algorithm. Generally, a stack is formed by a series of features differing from each other by the same *m/z* gap (or multiples of it), specifically those identified as common contaminating ions and adducts in ESI, http://www.waters.com/webassets/cms/support/docs/bckgrnd_ion_mstr_lst_4_13_2010.pdf (Keller, et al., 2008). Here, a lipid stack is a cluster of features with the same retention time. On the other hand, a contaminant stack comprises a series of features that elute with a gap between retention times that is maintained across the series. Thus, if plotting features in a *m/z* versus retention time scatter plot, lipid stacks are visualized as vertical sets whilst contaminant stacks are observed as diagonally spaced out features. Importantly, some lipid classes appear visually in the aforementioned scatter plots as clusters eluting in a diagonal fashion. For example, triglycerides that differ by 2C fragments and saturation/unsaturation follow predictable patterns in terms of retention time shift as the mobile phase lipophilicity increases. To avoid removing true lipids that follow this pattern, we have added the condition that a stack must have at least 4 features to be categorized as such before it is removed.

##### 2.2.3 Isotope removal

Isotope peaks due to the presence of one or more ^13^C atoms per molecule demonstrate a predictable pattern of features in scatter plots, eluting at the same retention time. These isotopes need to be either removed or combined with the molecular ion prior to searching databases for matches. This is important since related lipids often show multiple M+2 molecular species (differences in double bonds), and depending on instrument resolution an isotope with two ^13^C atoms could appear isobaric with a similar lipid to the molecular ion, but with one double bond less.

XCMS includes in its latest versions the CAMERA package which, among other features, detects and labels isotopic peaks. However, we found that the intensity ratio check used in its algorithm is not adequate for lipidomics, because the decrease in intensity of isotope peaks follows a non-linear relationship with the *m/z* increment in lipids (Yergey, 1983). Also, CAMERA labels a feature as an isotope if at least a chosen number of samples comply with the requirements (50% by default). Many studies in lipidomics involve diverse sample conditions, so different lipids (and isotopes) may be present only in some samples. Thus, CAMERA’s approach can be fallible, particularly since it does not distinguish between technical and biological replicates. We have developed a new deisotoping filter that calculates the isotopic distribution based on polynomial expansion of the parent intensity rather than the linear functions used in CAMERA. Furthermore, the conditions are checked independently for each set of technical replicates belonging to the same biological sample, treating them as separate entities. The method is described in detail in Appendix 2, and we have validated our formulae using the Yergey algorithm.

##### 2.2.4 Salt cluster removal

We have also implemented the salt cluster filter published by (McMillan, et al., 2016). This algorithm targets the common artefacts of ESI recognized by their particular high mass defects and early elution in LC/MS data. The method uses a mass-defect cutoff and a whitelist of bonafide metabolites (with high mass defects) to screen out the salt clusters.

##### 2.2.5 False Discovery Rate

The presence of diverse isotope and adduct features in samples is a critical challenge for the identification process, leading to false positives. The false discovery rate (FDR) is a common metric used to estimate the error of metabolite-spectrum matches and, hence, evaluate the confidence of the resulting profile (Schrimpe-Rutledge, et al., 2016). Although there is currently no agreed upon method to determine FDR in lipidomics, it is widely accepted in MS-based proteomics to use a target-decoy strategy (Elias and Gygi, 2007; Kall, et al., 2008).

The last feature added to *PeakFilter* is a FDR method developed adopting a similar approach. This uses target and decoy databases to measure robustness of a method, retrieving the number of hits from a dataset found in the decoy database (*n*_*d*_) versus the number of hits in the target database (*n*_*t*_), and calculating FDR as *n*_*d*_/*n*_*t*_. The target database is based on real molecules, while the decoy database is made up of false ions, e.g. *m/z* values which would not be expected in nature (described in detail below). In this method, the number of decoy hits would be considered to mirror the frequency of false lipid matches by LipidFinder’s putative profiling. Overall, FDR estimates provide a sense of robustness of the data after *PeakFilter*, that is, a theoretical estimate of how many lipid-like artefacts and/or non-lipid metabolites might remain afterwards.

To create its decoy database, JUMPm, a metabolite identification tool, altered the metabolites *m/z* values from the target database by violating the octet rule on chemistry (Jones, 2016). This means that each *m/z* value was adjusted to another theoretical value that did not fit this rule. We found that this *m/z* alteration does not work in lipidomics since many lipids differ from each other by the mass of one hydrogen, resulting in a decoy database that contains a significant percentage of *m/z* matching real lipids. Thus, we instead built a decoy database by adding 0.5 Da to every analyte, since this is a very rare mass defect in lipids. For our target database, we used an *in-silico* database (COMP_DB) composed of major classes of lipid species, generated from a list of commonly occurring acyl/alkyl chains. The almost 30,000 lipid species in COMP_DB are listed in “bulk format”, specifying total number of carbons and double-bonds in their constituent chains. Our decoy (COMB_DB_0.5Da) was generated from COMP_DB, as above. We show a validation and also an application of the approach using real data in Sections 4.1 and 4.4.

##### MS Search

Lipidomics datasets are large and complex, and there are significant challenges relating to both their identification and their subsequent visualisation. MS data can only ever provide bulk information on putative structures (e.g. potential lipid category, number of carbons and rings/double bonds on fatty acyl substituents, etc.). It cannot be used to define a lipid down to actual fatty acids, confirm head groups, assign either geometric isomers/enantiomers or position of functional groups (e.g. for oxidized groups). With this in mind, *m/z* searches must only return the correct bulk structures for MS data without providing detailed structural annotation. To address this issue, once lipidomics data has been processed through *PeakFilter*’s modules (Supplementary Figure 1) and amalgamated (if needed), *MS Search* performs putative mass assignment through a bulk structure search on LIPID MAPS. This module queries the selected repository lipid database to determine whether each remaining feature is a known analyte or not based on its *m/z*. This revised stage offers 3 in-house lipidomics databases:

**COMP_DB:** a computationally generated lipid database composed of about 30,000 bulk (isobaric) species covering the major classes of lipid species. It has been customized for precursor ion searching.
**ALL_LMSD:** the entire LIPID MAPS structure database (LMSD) of over 43,000 unique biologically relevant lipid structures.
**CURATED_LMSD:** a subset of approximately 21,000 curated structures from LMSD that have been reported in the literature (excluding computationally generated structures). *MS Search* allows the user specify mass tolerance and restrict the list of adducts and lipid categories being searched. The putative profile will only include matched lipids for a feature when searching against one of the LMSD databases and no bulk structures are found. The output file format has been changed from CSV to XLSX to enable inclusion of interactive content (via hyperlinks) for output results, as we did on LipidFinder’s online version (Fahy, et al., 2019).

Once putative matches have been assigned, visualization of data is required in a manner that allows the user to interpret the large amount of resulting data. This is a major challenge when dealing with often thousands of individual *m/z* values, including a large proportion of ions that have not been matched to any database entry. To aid this, *MS Search* now allows the user to create a summary file and a lipid category scatter plot from the putative identifications. The summary file retains information on the closest match of the most frequent lipid category for each unique *m/z* and retention time. To generate the lipid category scatter plot, the eight main lipid categories described by the LIPID MAPS Lipid Classification System (http://www.lipidmaps.org/data/classification/LM_classification_exp.php) are used, with the addition of an “unknown” category for unidentified features. Each lipid category is always assigned the same color, easing the comparisons between plots during the analysis. Additionally, LipidFinder 2.0 provides the option to use a color blind friendly palette.

It is important to note that unlike metabolites, lipids generally behave similarly across individual categories/classes in large cohort sample sets (e.g. phospholipids as a group tend to change together with disease or genotype), and it is highly unusual that a single lipid will stratify without others that are structurally related. We propose that taking the approach of plotting and statistically analyzing lipids in their LIPID MAPS categories has several advantages: **i)** if several lipids within a category alter similarly, it provides confidence that the difference is real, and **ii)** if analyzed in sub-groups, the statistical power is increased when applying corrections for multiple comparison testing. Our suggested approach is to screen categories for differences first using an untargeted method as described herein, then to validate the changes across the individual lipid categories using gold standard targeted quantitative MS/MS methods (generally restricted to single categories only) as the second step.

### Supplementary Results

#### FDR analysis

Decoy databases are designed to be populated with ions that should not commonly exist in the types of samples being analysed, and as expected, ours (COMP_DB_0.5Da) does not share any *m/z* values with COMP_DB ions. Thus, high quality lipidomics data from well-calibrated instruments, that has been effectively cleaned up should return a low FDR using a decoy-*versus*-target database approach, as only implausible lipids will be recognised by the decoy. To validate this approach, we created two “theoretical” sets of experimental data for testing, each of which contain 400 high resolution *m/z* values randomly selected from the target database, COMP_DB. The first is comprised of only [M−H]^−^ and [M+H]^+^ ions calculated from the neutral mass of real lipids (**molecular ions**) whilst the second includes several common lipid adducts: [M−H]^−^, [M+OAc]^−^, [M+H]^+^, [M+Na]^+^ and [M+NH_4_]^+^ (**molecular ions+adducts**). In both sets of ions, there are 200 of each polarity. See Appendix 4 for the detailed list of each dataset. To fully evaluate this approach, we performed the FDR test under two different database search conditions: **a)** limit the search to common lipid adducts: [M*−*H]^−^, [M+OAc]^−^, [M+H]^+^, [M+Na]^+^ and [M+NH_4_]^+^ (**common**) which are more representative of real lipidomics datasets, or **b)** screen for all available ions in the LIPID MAPS MS search tool, including a number of potential adducts of less direct relevance to lipids (**all**). We chose a standard mass tolerance for high resolution MS (± 0.001 Da) for both tests to fit with the expected instrumentation imperfections, e.g. suboptimal calibration, and other random electronic noise that will make real experiment *m/z* values differ slightly from the theoretical ones. Supplementary Table 1 shows the results for both datasets in each search condition. The number of features that returned at least one match in the target (*n*_*t*_) and in the decoy (*n*_*d*_) databases is also shown for completeness.

Supplementary Table 1 shows that our method returns a 0% FDR (with 100% *n*_*t*_) when searching for common lipid ions. These results demonstrate that the construction of a decoy database by adding 0.5 Da to every bulk lipid structure included in the target database is an effective strategy, since the method did not return any decoy hits even with a standard tolerance. Even if we expand our database search space to include every possible adduct available in the LIPID MAPS MS search tool, the FDR is negligible. This suggests that our proposed FDR approach can be used as a metric to assess the quality of LC/MS lipidomics datasets, e.g. retained artefactual ions or potential calibration drifts.

**Supplementary Table 1.**
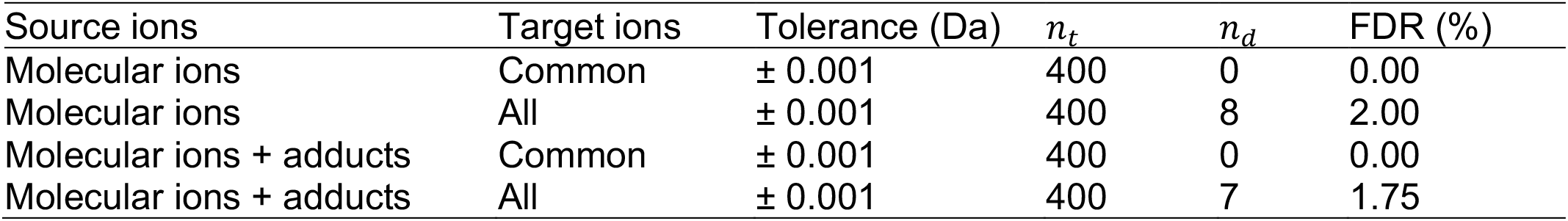
FDR of two sets of 400 ions calculated from randomly selected neutral lipid *m/z* of the target database (COMP_DB) and two different database search conditions. *n*_*t*_: number of target database matches; *n*_*d*_: number of decoy database matches.

##### Isotope removal analysis

Here we compared annotation of isotope peaks as achieved by CAMERA and LipidFinder 2.0 using a dataset of lipids generated by high-resolution LC/MS analysis of lipid extracts from RAW cells and peritoneal macrophages. Both approaches were applied to the datasets first pre-processed with XCMS Online with the Orbitrap II parameter set (Tautenhahn, et al., 2012). Following this, we ran *MS Search* to generate putative identifications of as many features as possible. This profiling was done searching against the COMP_DB database with a mass tolerance of ± 0.001 Da, and screening for all available ions (of the corresponding polarity). Supplementary Table 2 presents the isotope annotation returned by both software tools for 8 features (4 positive, 4 negative ions) and their putative identification.

**Supplementary Table 2.** Comparison of the isotope annotation of CAMERA *vs* LipidFinder 2.0 for 8 features, 4 in positive mode and 4 in negative mode, of the macrophages dataset right after being pre-processed with XCMS and putatively identified with *MS Search* (COMP_DB database).

| *m/z* | RT<br>(min) | Putative IDs |  | Software | Isotope | *m/z* | RT<br>(min) | Putative IDs |  |
| --- | --- | --- | --- | --- | --- | --- | --- | --- | --- |
|  |  | Bulk structure | Adduct |  |  |  |  | Bulk structure | Adduct |
| **668.6188** | 41.26 | DG(38:1) or | [M+NH<sub>4</sub>]<sup>+</sup> | CAMERA | M+1 | 669.6222 | 41.25 | unknown |  |
|  |  | TG(P-38:0) |  |  | M+2 | 670.6254 | 41.24 | TG(87:7) | [M+2H]<sup>2+</sup> |
|  |  |  |  | LipidFinder | M+1 | 669.6222 | 41.25 | unknown |  |
|  |  |  |  |  | M+2 | 670.6254 | 41.24 | TG(87:7) | [M+2H]<sup>2+</sup> |
| **831.6834** | 54.86 | PA(45:0) | [M+H]<sup>+</sup> | CAMERA | M+1 | 832.6873 | 54.85 | MGDG(38:0) | [M+NH<sub>4</sub>]<sup>+</sup> |
|  |  | DG(50:6) | [M+Na]<sup>+</sup> | LipidFinder | none |  |  |  |  |
| **878.8170** | 51.66 | TG(52:1) | [M+NH<sub>4</sub>]<sup>+</sup> | CAMERA | none |  |  |  |  |
|  |  |  |  | LipidFinder | M+1 | 879.8201 | 51.63 | unknown |  |
| **941.6996** | 52.08 | PA(54:8) | [M+H]<sup>+</sup> | CAMERA | M+1 | 942.7038 | 52.08 | MGDG(47:8) | [M+NH<sub>4</sub>]<sup>+</sup> |
|  |  | TG(56:8) | [M+K]<sup>+</sup> | LipidFinder | M+1 (Peritoneal) | 942.7038 | 52.08 | MGDG(47:8) | [M+NH<sub>4</sub>]<sup>+</sup> |
| **704.5594** | 40.27 | PE(O-34:0) | [M-H]<sup>-</sup> | CAMERA | none |  |  |  |  |
|  |  | PC(O-32:0) | [M-CH<sub>3</sub>]<sup>-</sup> | LipidFinder | M+1 | 705.5628 | 40.23 | unknown |  |
| **730.5391** | 40.21 | PE(35:1) | [M-H]<sup>-</sup> | CAMERA | M+1 | 731.5425 | 40.21 | unknown |  |
|  |  | PC(33:1) | [M-CH<sub>3</sub>]<sup>-</sup> | LipidFinder | M+1 (RAW) | 731.5425 | 40.21 | unknown |  |
| **751.5724** | 41.20 | unknown |  | CAMERA | M+1 | 752.5611 | 41.24 | PC(32:0) | [M+F]<sup>-</sup> |
|  |  |  |  |  | M+2 | 753.5590 | 41.23 | unknown |  |
|  |  |  |  | LipidFinder | none |  |  |  |  |
| **802.5756** | 41.41 | PE(P-42:6) | [M-H]<sup>-</sup> | CAMERA | M+1 | 803.5790 | 41.42 | unknown |  |
|  |  | PC(P-40:6) | [M-CH<sub>3</sub>]<sup>-</sup> | LipidFinder | M+1 | 803.5790 | 41.42 | unknown |  |

RT: retention time. The putative profile shows a few examples of the classes and adducts of the closest matches (smallest Δppm) with ±0.001 Da tolerance. If LipidFinder 2.0 annotated a feature as an isotope in only one of the biological sample groups (RAW or peritoneal) it is marked as so in the “Isotope” column.

Overall, CAMERA and LipidFinder 2.0 agreed on many isotope annotations. In several cases this agreement was over all sample replicates (6 RAW 264.7 and 6 resident peritoneal macrophages). For instance, in Supplementary Table 2 we find *m/z* 668.6188 (positive mode) and *m/z* 802.5756 (negative mode). On average, disagreement between both methods happened in less than 50% of replicates. As a result, CAMERA with its “majority rule” annotated whole features as isotopes that LipidFinder 2.0, with its more restrictive intensity thresholds, only annotated partially, i.e. defining those features as isotopes for some but not all sample replicates. This problem becomes critical when the discrepancy delineates the diverse biological groups of the dataset. See two examples in Table 2: *m/z* 941.6996 (+ve mode), whose M+1 isotope was found only in the peritoneal group; and *m/z* 730.5391 (-ve mode), whose M+1 isotope was only annotated in raw samples. In both cases the intensity was too low in the untagged group, meaning that those features are likely to be isotopes in their annotated group, and another type of analyte in the other.

Finally, we focus on features where each method has returned a different annotation. In Supplementary Table 2 we show *m/z* 831.6834 (+ve mode) and *m/z* 751.5724 (-ve mode), for which LipidFinder 2.0 did not detect isotopes in any sample replicate whilst CAMERA did. In the former instance, the isotope has a higher intensity than our defined upper threshold for every sample. In the negative example, the mass of the feature annotated as the M+1 isotope by CAMERA is much lower than the expected increment of one ^13^C: the absolute difference between the “parent” and its isotope is 0.9887 Da (instead of 1.0033 Da). Furthermore, both M+1 isotopes were putatively identified as MGDG(38:0) and PC(32:0), respectively, another reason to believe they may have been incorrectly annotated as isotopes. Conversely, Supplementary Table 2 contains *m/z* 878.8170 (+ve mode) and *m/z* 704.5594 (-ve mode) as examples where LipidFinder detected isotopes in every sample replicate whilst CAMERA did not. The latter method is assigning different peak groups (pcgroup) to the “parent” features and their corresponding isotopes, even though they are less than 3 seconds apart from each other in both cases.

##### Overall performance analysis

LipidFinder 2.0 includes several new modules all designed to improve data quality. To demonstrate this, we provide a comparison of effectiveness and performance of the lipidomics pipeline (Supplementary Figure 2) with and without either the old or new versions of LipidFinder. For XCMS we chose the online version with the Orbitrap II set of parameters. We used the output files from XCMS Online as input for both LipidFinder versions, with the default parameterization of LipidFinder 2.0 applied in both cases. Supplementary Figure 2 shows the retained features of the macrophages dataset in both positive and negative mode in scatter plots of *m/z* versus retention time. Last, an example of the color blind pallet for both a scatter plot and volcano plot of the data is presented in Supplementary Figures 3,4.

Prior to using LipidFinder, XCMS datasets contain eluting features that form repeating patterns (Supplementary Figure 2, left panels). For example, series of dots following (almost) straight lines. These are in many cases related to well-known artefacts such as in-source fragments, lipid stacks, ESI contaminants or salt clusters. Applying the FDR method to these XCMS datasets we obtain high %FDR values, indicating that many features match our decoy lipid *m/z* values (Supplementary Table 3). LipidFinder 1.0 performs reasonably at removing several of these artefacts, resulting in a reduced FDR (Supplementary Table 3, Supplementary Figure 2, middle panels). However, significant artefacts still remain, e.g. the dense column of dots throughout the first minute (salt clusters), or the short vertical lines of equally distant dots (isotopes). In contrast, the incorporation of the new modules presented here into the lipidomics pipeline results in further improvements to data output as shown by improved FDR (Supplementary Table 3, Supplementary Figure 2, right panels). The first release of LipidFinder removed a third of the total features in each dataset and reduced the number of decoy hits up to a fourth of that of XCMS Online. The reduction of target hits is also expected as LipidFinder has modules designed to “simplify” the dataset. For instance, adduct removal will find all adducts for a given lipid and retain only the most intense one. LipidFinder 2.0 improves on the results of its predecessor removing two thirds of the total number of features from XCMS Online, but retaining almost as many target hits as LipidFinder 1.0 and decreasing FDR by half. Here, the minor reduction of target hits is due mainly to in-source fragment removal, e.g. where fatty acids will be retained only if they do not co-elute with other features from which they may have originated.

**Supplementary Table 3.** FDR of macrophages datasets (both polarity modes) after XCMS Online, either alone or followed by LipidFinder 1.0 or 2.0. *n*: total number of features; *n*_*t*_: number of target database matches; *n*_*d*_: number of decoy database matches.

| Processing | Polarity | $n$ | $n_t$ | $n_d$ | FDR (%) |
| --- | --- | --- | --- | --- | --- |
| XCMS Online | Negative | 6336 | 2020 | 214 | 10.59 |
| +LipidFinder 1.0 | Negative | 4222 | 1000 | 69 | 6.90 |
| +LipidFinder 2.0 | Negative | 2350 | 937 | 37 | 3.95 |
| XCMS Online | Positive | 16611 | 5792 | 1308 | 22.58 |
| +LipidFinder 1.0 | Positive | 10671 | 2553 | 318 | 12.46 |
| +LipidFinder 2.0 | Positive | 5387 | 2451 | 193 | 7.87 |

Our previous version of LipidFinder typically took 30 minutes to analyze 12 samples (with about 23K features for each mode). In the new version, we sought to improve processing speed through parallelization and algorithm redesign based on computational cost analysis. Here we compare time taken to process the macrophages datasets (positive and negative mode) after using the same configurations and conditions as for the previous analysis, using both versions and finding that the total has reduced by 10 minutes (Supplementary Table 4). The benchmarks show increased time cost for *PeakFilter*. This is due to the computational requirements of the FDR method, creating a bottleneck. If we omit that step, the new implementation outperforms its predecessor, even with the addition of three new modules. We observe similar improvements in *Amalgamator* and *MS Search*. Furthermore, during the algorithm redesign aforementioned we have paid special attention to the computational complexity. This means that not only is LipidFinder 2.0 more efficient, but its time cost will increase more slowly with larger datasets.

##### LipidFinder on LIPID MAPS

LipidFinder 2.0 replaces LipidFinder 1.0 on its online version, available on LIPID MAPS as a web application (Fahy et al., 2018). As an additional feature, LipidFinder on LIPID MAPS incorporates the option to import pre-processed files directly from XCMS Online. To do so, users are requested their username from XCMS Online and the JobID that has generated the pre-processed file they want to import. This feature will only fetch datasets from pairwise and multigroup job types. After the file has been imported correctly, it can be processed with LipidFinder 2.0 and analysed with the statistical methods available.

**Supplementary Table 4.** Benchmark results of LipidFinder’s old and new implementations for the same macrophage datasets and parameters.*PeakFilter* version 2.0 includes all the new additional modules. The time in parenthesis corresponds to the time cost without the FDR module. Selected CURATED_LMSD database to compare *Web Search* (v1.0) and *MS Search* (v2.0).

| Version | *PeakFilter* |  | *Amalgamator* | *MS Search* |
| --- | --- | --- | --- | --- |
|  | Negative mode | Positive mode |  |  |
| 1.0 | 59s | 163s | 21s | 1539s |
| 2.0 | 122s (38s) | 322s (95s) | 12s | 1261s |

**Supplementary Figure 2.**
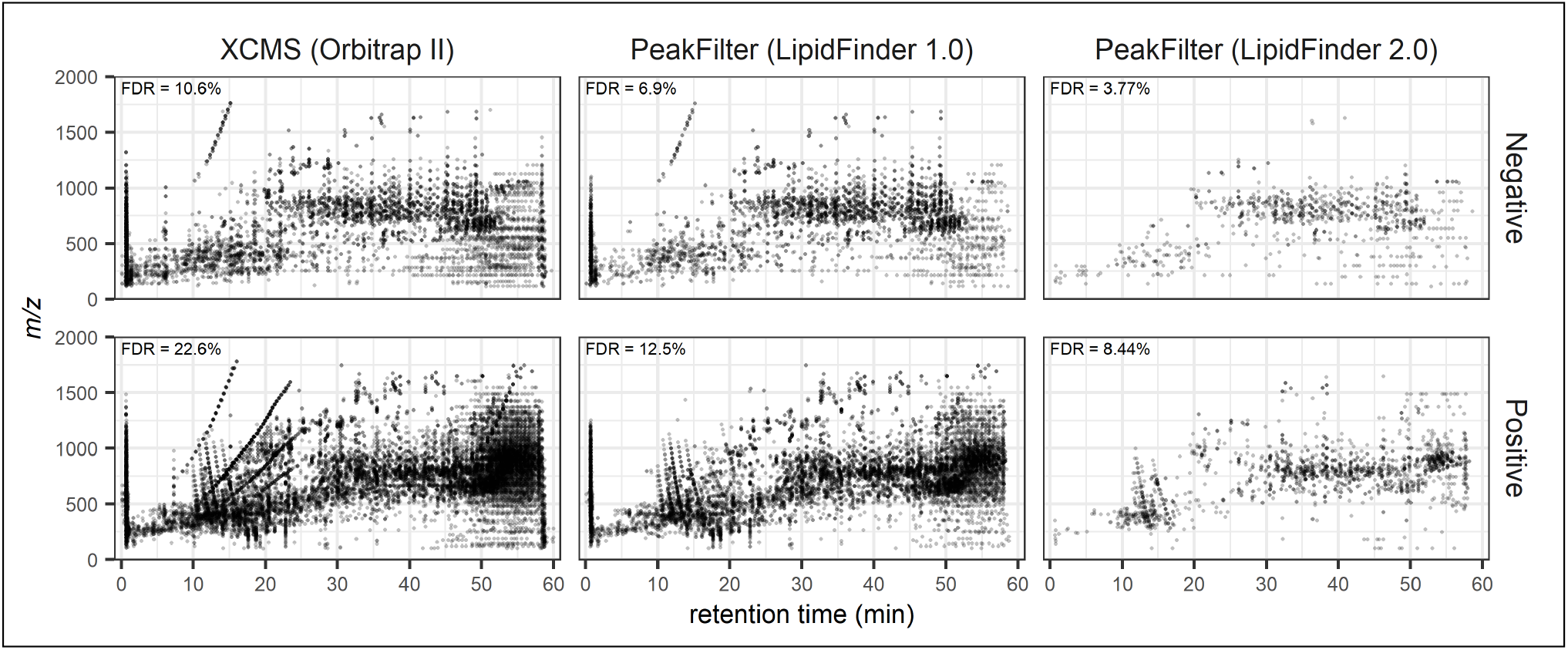
Comparison of the number of features retained after XCMS, and after *PeakFilter* for LipidFinder 1.0 and LipidFinder 2.0. These results are based on the macrophages dataset (RAW and peritoneal) in both polarity modes. Used XCMS Online with Orbitrap II parameters, and the default parameter values of LipidFinder 2.0 were chosen for both versions of LipidFinder. The FDR is included in every

**Supplementary Figure 3.**
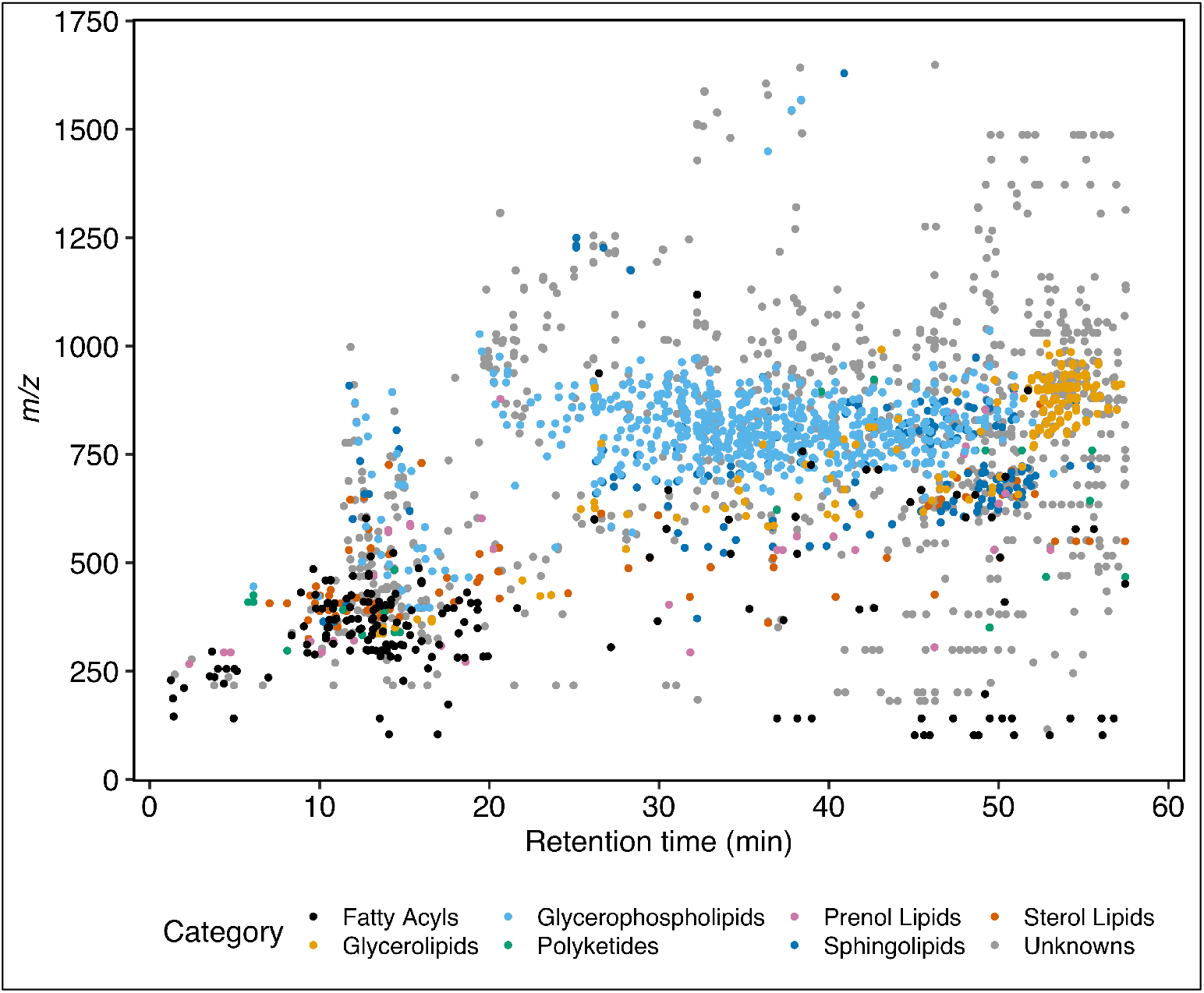
Example of lipid category scatter plot for macrophages data. Both polarity modes run through LipidFinder 2.0 with the default parameters and amalgamated. *MS Search* performed against COMP_DB database with 0.001 Da tolerance.

**Supplementary Figure 4.**
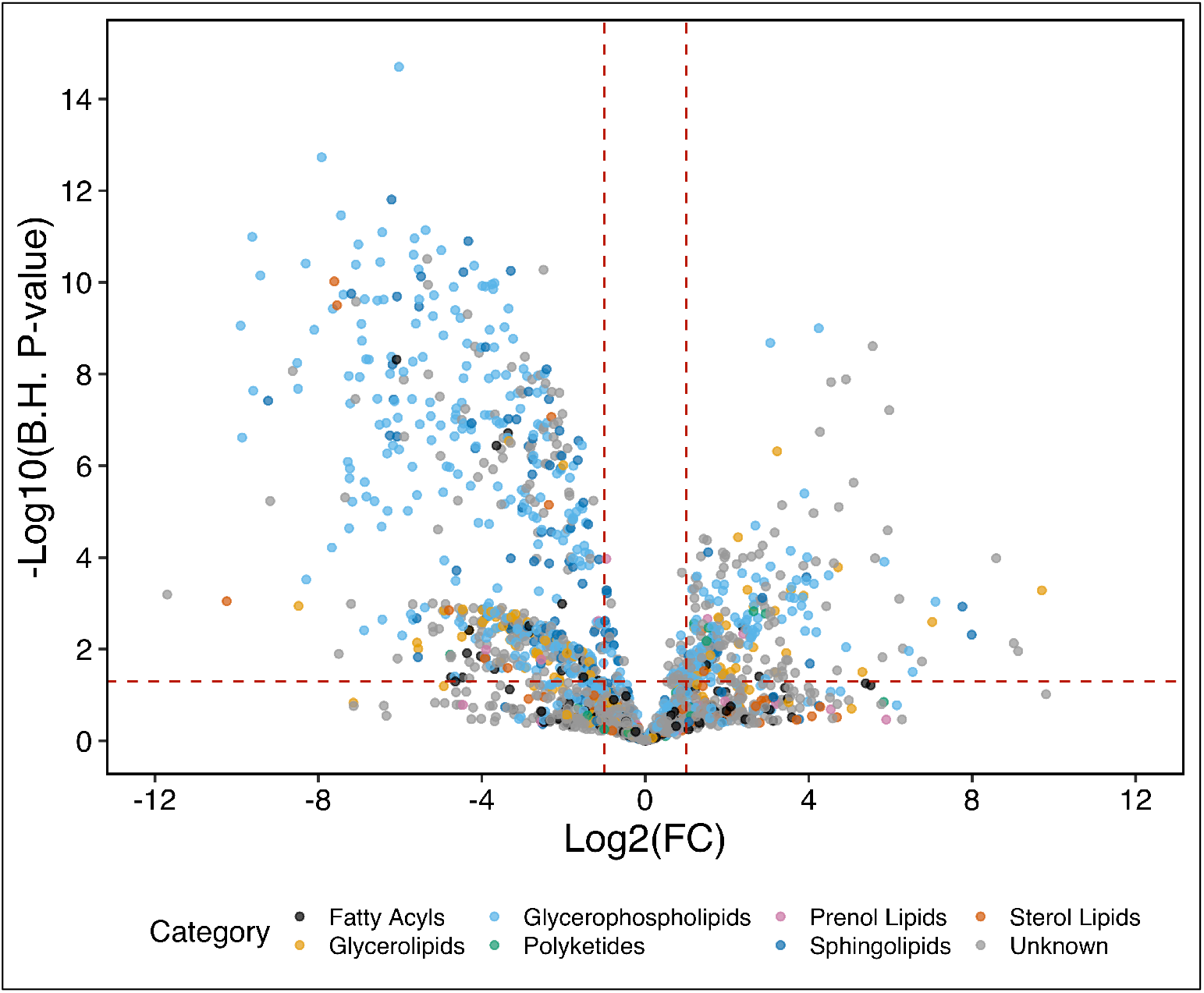
Volcano plot of significantly increased lipids in both pResMϕ and RAW 264.7 cells by lipid category. Mϕ lipid extracts from RAW cells and pResMϕ were analyzed by LC-FTMS on the Orbitrap Elite, at 60,000 resolution, then processed using XCMS followed by LipidFinder. Volcano plots representing fold-change and p-value of lipids in pResMϕ relative RAW cells by lipid category. Statistical significance was determined using the Holm-Sidak method, with alpha=5.00%. Each row was analysed individually, without assuming a consistent SD. Differential expression was classified as significant when P < 0.05 and |FC| > 1.5 (depicted in black). Both polarity modes run through LipidFinder 2.0 with the default parameters and amalgamated. *MS Search* performed against COMP_DB database with 0.001 Da tolerance.

## Appendix 1. In-source ion fragmentation removal

Default list of in-source fragments and common neutral losses. In all cases the parent *m/z* has to be above 400:

- Common phospholipid (PLs) fragments (*m/z* values) to remove from the dataset in negative polarity mode: 237.4042, 255.233, 261.4229, 263.4388, 265.4546, 279.233, 281.2486, 283.2643, 285.4444, 303.233, 309.4659, 311.4187, 327.233, 329.2486.
- Neutral *m/z* loss for fatty acids (FA) in both negative and positive polarity modes: 256.2402, 280.2402, 282.2559, 284.2715, 304.2402, 328.2402, 330.2559.
- Neutral *m/z* loss for ketenes in both negative and positive polarity modes: 238.4094, 262.4308, 264.4467, 266.4626, 286.4523, 310.4738, 312.4896.
- Neutral *m/z* loss for phosphatidylserines (PS) in negative polarity mode: 78.9591, 87.0321, 96.9696, 152.9958.
- Neutral *m/z* loss of headgroup and a fatty acid for PS in negative polarity mode: 326.4948, 344.3256, 350.5162, 352.5321, 354.548, 368.3256, 370.3413, 372.3569, 374.5377, 392.3256, 398.5592, 400.575, 416.3256, 418.3413
- Neutral *m/z* loss for PS in positive polarity mode: 87.0321, 185.0727.
- Neutral *m/z* loss for phosphatidic acids (PA) in negative polarity mode: 78.9591, 96.9696, 152.9958.
- Neutral *m/z* loss for PA in positive polarity mode: 97.9952.
- Neutral *m/z* loss for phosphatidylethanolamines (PE) in positive polarity mode: 141.0631.
- Neutral *m/z* loss for phosphatidylglycerols (PG) in positive polarity mode: 172.0739.
- Neutral *m/z* loss for phosphatidylinositols (PI) in negative polarity mode: 78.9591, 96.9696, 152.9958, 241.1128.
- Neutral *m/z* loss for PI in positive polarity mode: 260.1360.
- Neutral *m/z* loss of H_2_O in both negative and positive polarity modes: 18.0153.
- Neutral *m/z* loss of CO_2_ in both negative and positive polarity modes: 44.0095.
- Neutral *m/z* loss of both H_2_O and CO_2_ in both negative and positive polarity modes: 62.0248.
- Neutral *m/z* loss of NH_3_ and a fatty acid for NH_4_^+^ adducts of diglycerides and triglycerides in positive polarity mode: 273.4552, 297.4767, 299.4926, 301.5084, 321.4981, 345.5196, 347.5355.
- Neutral *m/z* loss of H_2_O and NH_4_^+^ for diglycerides in positive polarity mode: 35.0458.

High-level description of the in-source fragmentation removal algorithm:

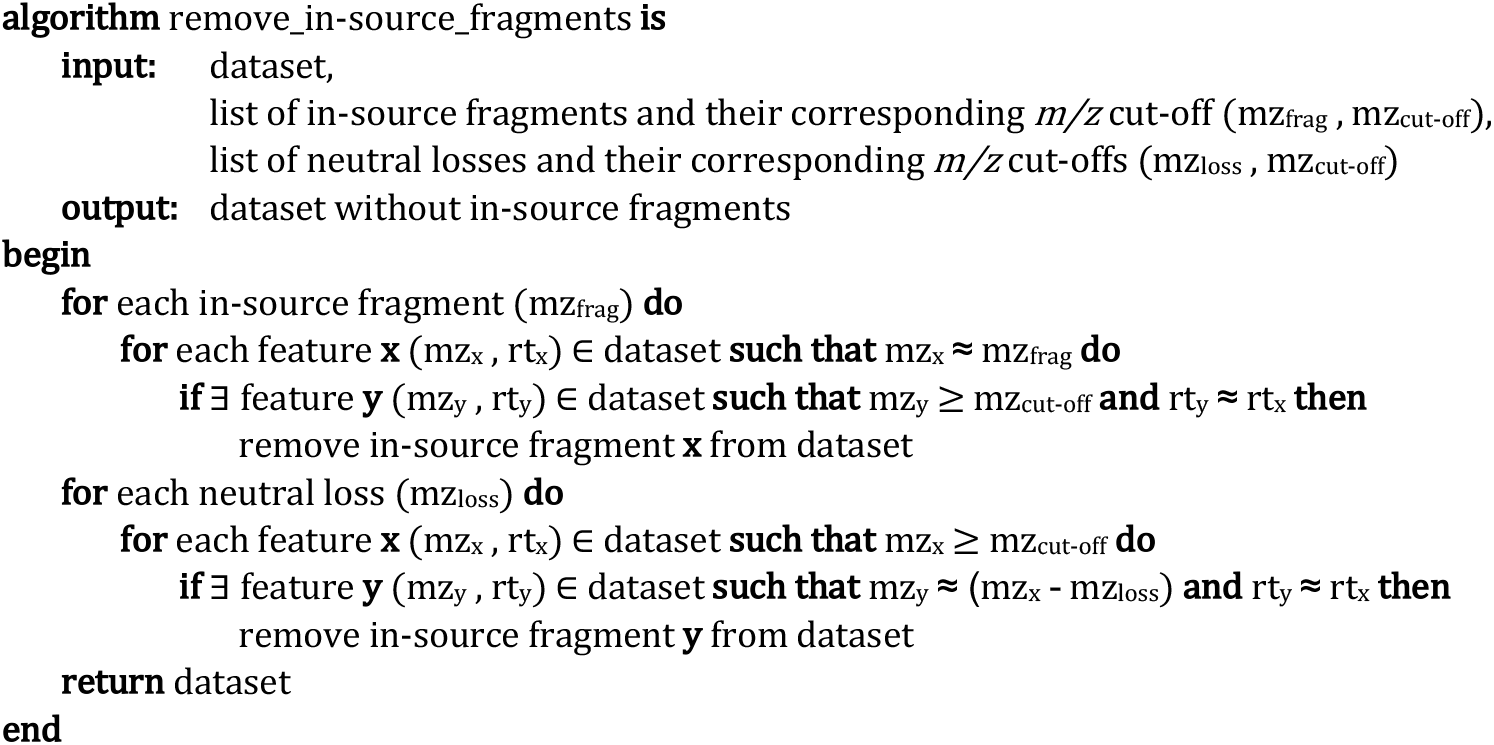

## Appendix 2. Isotope removal algorithm

CAMERA is used by XCMS to detect and annotate diverse ion species, including adducts and isotopes. However, its algorithm is not adequate to properly identify lipid isotopes because it uses a linear intensity ratio check to identify them, when this type of isotopic peaks has been shown to increase exponentially with their *m/z* value (Yergey, 1983). Moreover, the resultant thresholds are too broad, thus low-intensity lipids might be misidentified as isotopes. This is of special relevance for the first (M+1) and second (M+2) isotopes. We have implemented in LipidFinder an algorithm that replaces CAMERA’s intensity threshold equations for the first two isotopes (equations (1) and (2)) by the appropriate polynomial expansion ones (equations (3) and (4)). Following the same approach as CAMERA, we calculate a rough approximation of the number of carbons in a lipid as *numC* = ⌈*m/z*/12⌉.

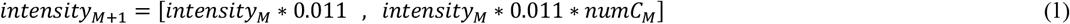

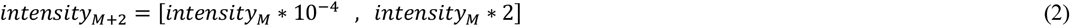

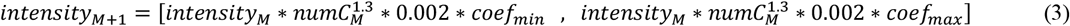

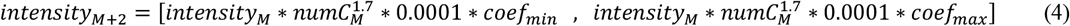

For illustration purposes, Supplementary Figure 5 shows the previously mentioned differences regarding intensity thresholds for isotopes M+1 and M+2 from a default parent intensity of 100 and diverse *m/z* values, applying LipidFinder’s default values for the minimum and maximum coefficients (0.7 and 1.3, respectively). As we can see, the tolerance in our algorithm is much narrower than that of CAMERA (especially for M+2) and it increases proportionally with the *m*/*z* of the parent in both cases.

**Supplementary Figure 5.**
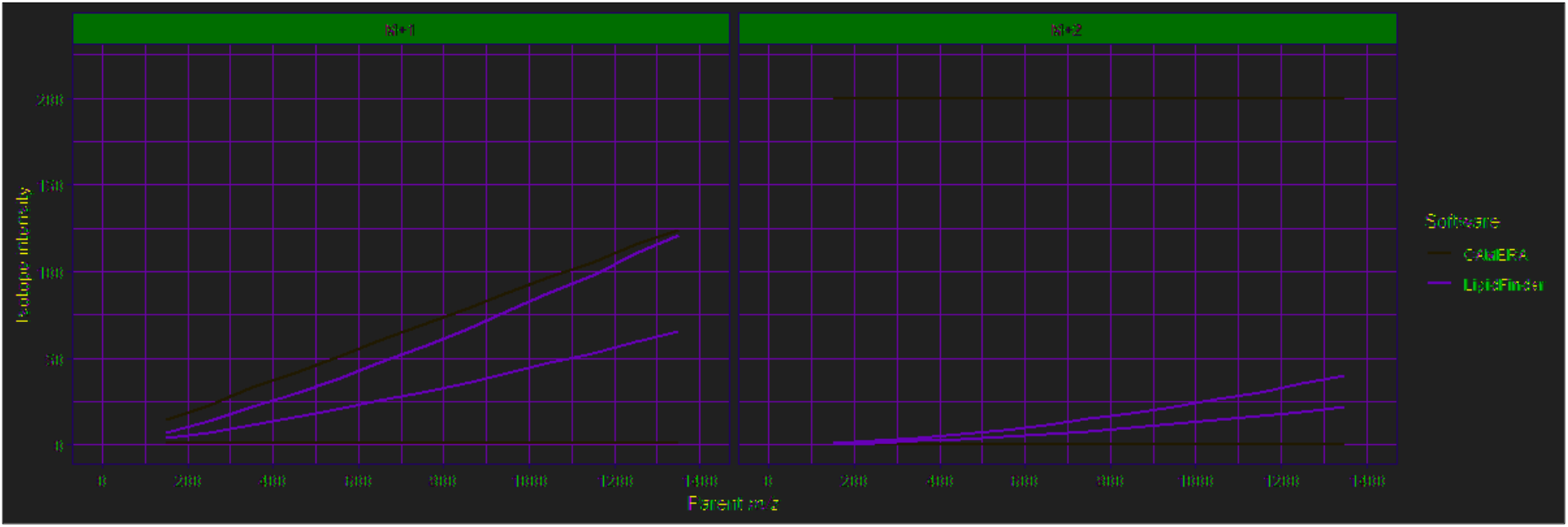
Comparison of M+1 and M+2 isotopes’ intensity thresholds for CAMERA and LipidFinder. The minimum and maximum intensity thresholds have been calculated from a parent *m/z* that varies from 150 to 1350 and a fixed parent intensity of 100. For LipidFinder, the minimum and maximum coefficients applied are 0.7 and 1.3, respectively.

As we indicate earlier, we have found cases where a feature matches the *m/z*, retention time and intensity conditions to be an isotope of another analyte, but it has not been annotated as such by CAMERA because its algorithm has assigned a different peak group (pcgroup) to it. To address this problem, LipidFinder’s method analyses every feature that is within the isotopic *m/z* and same retention time as the parent, and then applies the intensity thresholds to find the isotope. Although it is likely to be less efficient for large datasets, since the algorithm explores the whole dataset for every analyte, we consider this approach to be more accurate.

There is one last main difference between both methods: instead of annotating a feature as an isotope only if that feature meets the conditions in at least half of the samples (CAMERA’s default), our isotope removal algorithm is applied to each biological sample independently.

Finally, we include a high-level description of the isotope removal algorithm applied to each biological sample (note that C13 offset represents the exact difference between ^13^C and ^12^C isotopes):

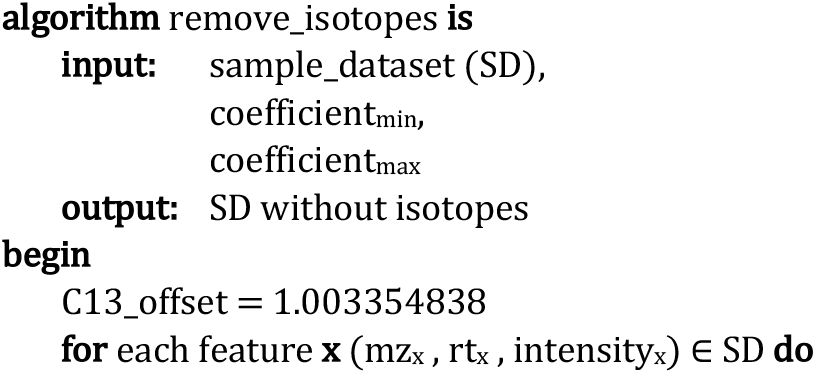
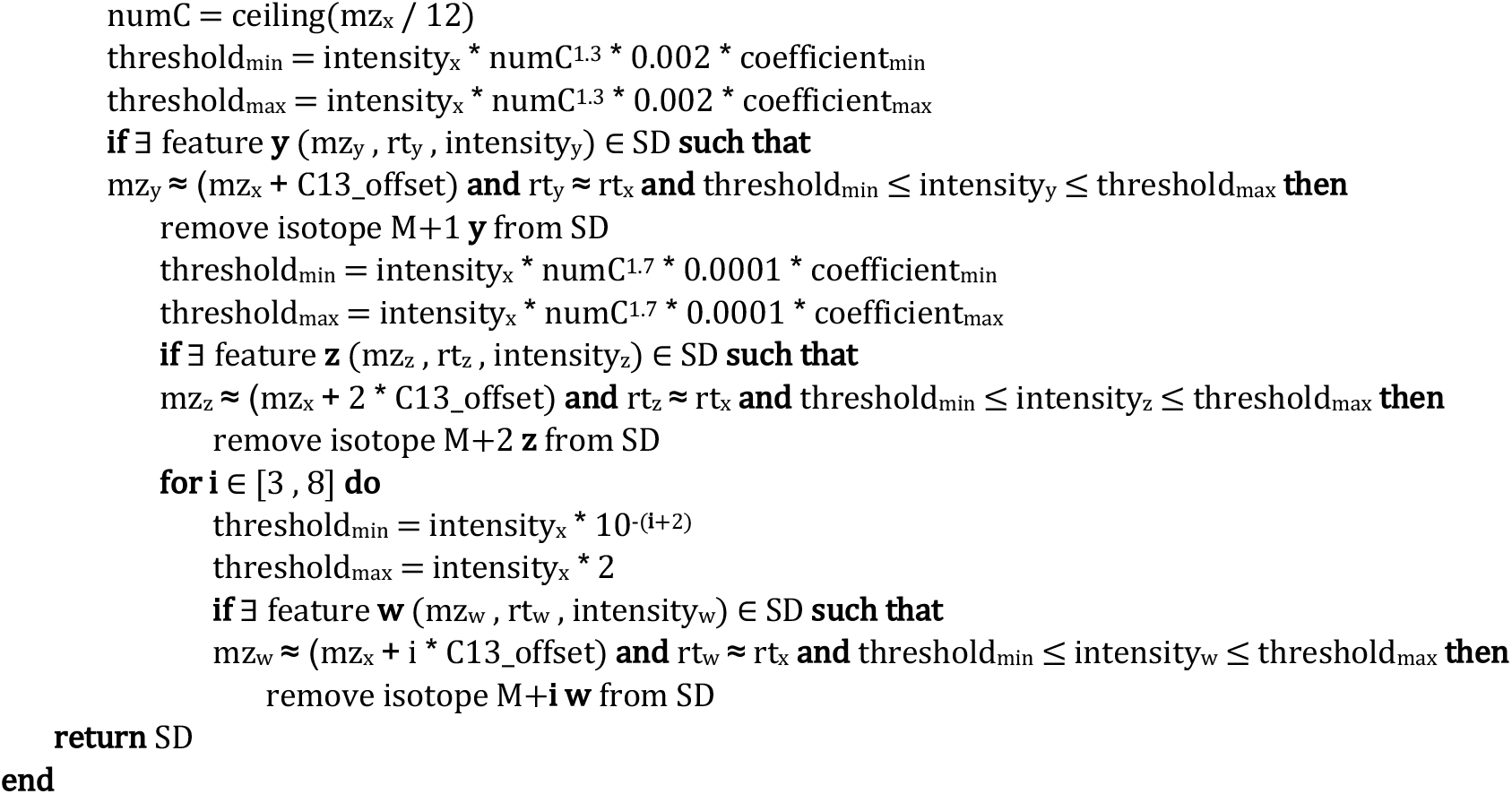

## Appendix 3. Macrophage dataset

The real-case macrophages dataset example was obtained in the laboratory as follows:

1. **Resident peritoneal macrophages** C57BL/6 mice were killed by asphyxiation using CO_2_, and death confirmed using cervical dislocation. Peritoneal cells were obtained by peritoneal lavage with 5 ml of PBS containing 5 mM EDTA and kept on ice until use. Resident macrophages were identified in all experiments as F4/80 ^+^ CD11b ^+^ CD73^+^ and MHCII^low^ cells and isolated by Fluorescence activated cell sorting.
2. **Culture of RAW 264.7** Murine macrophage-like RAW 264.7 cells were cultured in DMEM containing 0.26 mg/mL L-glutamine, 10% (v/v) fetal calf serum, 1 % Penicillin-Streptomycin (GE Healthcare, Cardiff, UK)). Cells were routinely grown in T175 flasks at 37 °C and 5 % CO_2_ and subcultured by trypsination at 1:3 ratio.
3. **Global lipidomics** Following harvesting of cells, lipids were extracted using two consecutive liquid-liquid extractions. First, hexane:isopropanol:acetic acid, then a modified Bligh and Dyer method (Bligh and Dyer, 1959), then resuspended in methanol and stored at −80 °C until LC/MS/MS. For this, lipids were extracted by adding solvent (1M acetic acid/propan-2-ol/hexane; 2:20:30; v/v/v) in a ratio of 2.5 mL solvent per 1 mL sample, and then vortexing for 1 min. Hexane (2.5 mL) was added, samples vortexed for 1 min and centrifuged for 5 min at 500 g, 4°C. The upper hexane layer was collected. The sample was re-extracted by adding hexane (2.5 mL) to the remaining aqueous phase, vortexing for 1 min and centrifuging for 5 min at 500 g, 4°C. Again, the upper hexane layer was collected and combined with the first hexane layer. The remaining aqueous phase was re-extracted as follows: 3.75 mL solvent (chloroform/methanol; 1:2; v/v) was added per sample. After vortexing for 1 min., 1.25 mL chloroform was added, and vortexed again for 30 sec., before adding 1.25 mL water, followed by 30 sec. vortex. Samples were centrifuged for 5 min. at 500 g and 4°C and the bottom chloroform layer collected and combined with the two hexane layers before drying under vacuum. Samples were re- suspended in 200 μL methanol. Lipid extracts were separated on an Accucore C18 column (150 × 2.1 mm, 2.6 μm) at a flow rate of 0.425 mL/min at 30 °C. All samples were loaded via Nexera X2 autosampler at 4 °C. The mobile phase consists of A (H2O:ACN 80:20 v/v) and a phase B (IPA: ACN, 70:30 v/v). The column was equilibrated in Solvent B 16%, and 20μL sample (dissolved in MeOH) were injected. A linear gradient was optimised as follows: 0 – 12 min, 16 – 60 % B; 12 – 19 min, 60 – 72 % B; 19 – 42 min, 72 – 84 % B; 42 – 51 min, 42 – 100 % B; and holding 100 % B for 6 min, followed by returning to solvent 16 % B and holding for 8 min for re-equilibration. MS conditions were as follows for analysis in positive ESI ionization mode: resolution 60,000 at 400 *m/z* HESI-II temperature 400 °C, N_2_ as drying gas, sheath gas flow 37 arbitrary units (au), auxiliary gas flow 15 au, sweep gas flow 1 au, capillary temperature 320 °C, spray voltage + 4.0 kV, S-lens RF level 62 %. Lock mass was *m/z* 391.2843. For analysis in negative ESI ionization mode: resolution 60,000 at 400 *m/z*, HESI-II temperature 350°C, N_2_ as drying gas, sheath gas flow 37 au, auxiliary gas flow 15 au, sweep gas flow 2 au, capillary temperature 320 °C, spray voltage – 3.5 kV, S-lens RF level 69 %. Lock mass was *m/z* 265.1479. A sample dataset was created by analyzing 6 technical replicates of peritoneal lavage extracts, and 6 technical replicates of RAW cell extracts, prepared as above, with 2 blanks of phosphate-buffered saline also included. The entire batch was run on the Orbitrap in both positive and negative mode, then processed together, first using XCMS Online (Orbitrap II settings), followed by LipidFinder 1.0 or 2.0 using LipidFinder 2.0 default parameter settings.

## Notes

### Competing Interest Statement

The authors have declared no competing interest.

## References

Bowden, J.A., et al. Harmonizing lipidomics: NIST interlaboratory comparison exercise for lipidomics using SRM 1950-Metabolites in Frozen Human Plasma. Journal of lipid research 2017;58(12):2275–2288.

Cappadona, S., et al. Current challenges in software solutions for mass spectrometry-based quantitative proteomics. Amino Acids 2012;43(3):1087–1108.

Fahy, E., et al. LipidFinder on LIPID MAPS: peak filtering, MS searching and statistical analysis for lipidomics. Bioinformatics 2019;35(4):685–687.

Liebisch, G., Ejsing, C.S. and Ekroos, K. Identification and Annotation of Lipid Species in Metabolomics Studies Need Improvement. Clin Chem 2015;61(12):1542–1544.

O'Connor, A., et al. LipidFinder: A computational workflow for discovery of lipids identifies eicosanoid-phosphoinositides in platelets. JCI Insight 2017;2(7):e91634.

O'Donnell, V.B., Murphy, R.C. and Watson, S.P. Platelet lipidomics: modern day perspective on lipid discovery and characterization in platelets. Circ Res 2014;114(7):1185–1203.

Wenk, M.R. The emerging field of lipidomics. Nat Rev Drug Discov 2005;4(7):594–610.

## References

Bligh, E.G. and Dyer, W.J. A rapid method of total lipid extraction and purification. Canadian journal of biochemistry and physiology 1959;37(8):911–917.

Covey, T., Lee, ED, Bruins, AP, Henion, JD. liquid chromatography/mass spectrometry. Analytical Chemistry 1986;58(14):1451A–1461A.

Elias, J.E. and Gygi, S.P. Target-decoy search strategy for increased confidence in large-scale protein identifications by mass spectrometry. Nat Methods 2007;4(3):207–214.

Gabelica, V. and De Pauw, E. Internal energy and fragmentation of ions produced in electrospray sources. Mass Spectrom Rev 2005;24(4):566–587.

Jones, D., Wang, X, Shaw, T, Cho, JH, Chen, PC, Dey, KK, Zhou, S, Li, Y, Kim, C, Taylor, JP, Kolli, U, Peng, J. Metabolome identification by systematic stable isotope labelling experiments and false discovery analysis with a target-decoy strategy. BioRxiv 2016.

Kall, L., et al. Assigning significance to peptides identified by tandem mass spectrometry using decoy databases. J Proteome Res 2008;7(1):29–34.

Keller, B.O., et al. Interferences and contaminants encountered in modern mass spectrometry. Anal Chim Acta 2008;627(1):71–81.

Lommen, A. and Kools, H.J. MetAlign 3.0: performance enhancement by efficient use of advances in computer hardware. Metabolomics 2012;8(4):719–726.

McMillan, A., et al. Post-acquisition filtering of salt cluster artefacts for LC-MS based human metabolomic studies. J Cheminform 2016;8(1):44.

Pluskal, T., et al. MZmine 2: modular framework for processing, visualizing, and analyzing mass spectrometry-based molecular profile data. BMC Bioinformatics 2010;11:395.

Schrimpe-Rutledge, A.C., et al. Untargeted Metabolomics Strategies-Challenges and Emerging Directions. J Am Soc Mass Spectrom 2016;27(12):1897–1905.

Slatter, D.A., et al. Mapping the Human Platelet Lipidome Reveals Cytosolic Phospholipase A2 as a Regulator of Mitochondrial Bioenergetics during Activation. Cell Metab 2016;23(5):930–944.

Smith, C.A., et al. XCMS: processing mass spectrometry data for metabolite profiling using nonlinear peak alignment, matching, and identification. Anal Chem 2006;78(3):779–787.

Smith, R., Ventura, D. and Prince, J.T. LC-MS alignment in theory and practice: a comprehensive algorithmic review. Brief Bioinform 2015;16(1):104–117.

Tautenhahn, R., et al. XCMS Online: a web-based platform to process untargeted metabolomic data. Anal Chem 2012;84(11):5035–5039.

Weckwerth, W. Metabolomics in systems biology. Annu Rev Plant Biol 2003;54:669–689.

Xu, Y.F., Lu, W. and Rabinowitz, J.D. Avoiding misannotation of in-source fragmentation products as cellular metabolites in liquid chromatography-mass spectrometry-based metabolomics. Anal Chem 2015;87(4):2273–2281.

Yergey, J., Heller, D, Hansen, G, Cotter, RJ, Fenselau, C. Isotopic distributions in mass spectra of large molecules. Analytical Chemistry 1983;55(2):353–356.

Zhang, W., et al. MET-COFEA: a liquid chromatography/mass spectrometry data processing platform for metabolite compound feature extraction and annotation. Anal Chem 2014;86(13):6245–6253.

Zhou, B., et al. LC-MS-based metabolomics. Mol Biosyst 2012;8(2):470–481.

